# The paradoxical relationship between CRISPR-Cas and phage susceptibility in *Pseudomonas aeruginosa*

**DOI:** 10.1101/2022.09.29.510058

**Authors:** Cédric Lood, Emma Verkinderen, Aleksandra Petrovic Fabijan, Jonathan R. Iredell, Pieter-Jan Ceyssens, Rob Lavigne, Vera van Noort

## Abstract

CRISPR-Cas systems are part of the pan-immune system of *Pseudomonas aeruginosa* and have been shown to limit horizontal gene transfers in that species. Indeed, isolates equipped with these systems tend to have smaller genomes and CRISPR spacers targeting integrative conjugative elements, phages, and plasmids. In this work, we investigate the genomic effects and phenotypic consequences of CRISPR-Cas systems in *P. aeruginosa*. First, we establish that the population structure is a confounding factor of the relation between genome sizes and the presence of CRISPR-Cas systems in *P. aeruginosa* as isolates from group II are, on average, 200 kbp larger than those from group I, and have a lower likelihood of possessing CRISPR-Cas systems. Second, we show that the impact on the genome size of CRISPR deactivation by anti-CRISPR proteins differs between the various CRISPR-Cas types found in this species (I-C, I-E, and I-F). Finally, we highlight a paradoxical, positive correlation between the presence of CRISPR-Cas systems and the chances of the host being infected by a set of distinct, strictly virulent Pseudomonas phages. We propose that this increased phage susceptibility in the presence of CRISPR-Cas is linked to a depletion of other accessory defense system genes in isolates with CRISPR-Cas systems.

<u>Note</u>: This manuscript was published as a chapter in the doctoral dissertation of Cédric Lood [72]

## Introduction

CRISPR-Cas systems are part of the adaptive and inheritable immune system of prokaryotes. The scientific story of the discovery of their genetic signature [1], the elucidation of their function as adaptive defense mechanisms against bacteriophages and mobile genetic elements [2]–[4], and later repurposing as a flexible genetic engineering tool in eukaryotes [5] is one that is both fascinating and abundantly documented [6]–[9]. These systems have been detected in multiple bacterial and archaeal species and appear to be accessory across those domains of life, with about 42% of the bacteria and 85% of archaea surveyed in 2020 by Makarova *et al*. equipped with such systems [10]. At the species level, it also appears that CRISPR-Cas is part of the accessory genome content of the population [11]–[14]. CRISPR-Cas systems share the feature of having CRISPR arrays that are populated with so-called spacer elements containing genetic signatures revealing the adaptive immunity. These systems are diverse both in evolutionary terms and in their mode of operation [15], and the latest classification scheme divides them into two classes, which are further split into six types and 17 subtypes [10].

The presence of these systems in the accessory genome content of various bacterial populations has raised interesting questions regarding their utility and cost for the organisms that carry them, and the coevolution dynamics between these systems and phages [16], [17]. For instance, bacteria coming under predation may either undergo constitutive defense selection (e.g., by masking a membrane receptor or through genomic mutations) or an inducible defense such as a CRISPR-Cas adaptive immune response. As both of these routes are observed in nature, it is likely that for a given species, the relative tradeoffs in costs between the two is determined by the specifics of the infection cycles of the bacteria and its parasites [18]. Specifically in the case of CRISPR, recent modelling efforts have shown that the immune response carries a cost that likely depends on both the regulation of the system and the timing of (or delay in) its effective response in clearing infections [19]. Adding to these considerations is the emerging observation that CRISPR-Cas systems shape the genomes of their hosts by restricting horizontal gene transfer [12], [20] and that the selective deactivation of CRISPR-Cas systems could be driving niche adaptation [21]. The shaping of genomes by CRISPR-Cas systems was shown recently for *P. aeruginosa* where (i) the distribution of genome sizes appears to be bimodal, (ii) the presence of CRISPR-Cas systems was linked to smaller genome sizes of the population carrying the system, and (iii) spacers were used to show evidence of genomic islands targeting [22].

The evolutionary dynamics between bacteria and phages is often usefully thought of as an arms race [23], and this is well illustrated by considering the case of CRISPR-Cas and anti-CRISPR [24]. Originally discovered in the prophages of *Pseudomonas* [25], the anti-CRISPR (Acr) proteins interfere with the functioning of CRISPR-Cas systems [26], [27]. These proteins are diverse and the current catalogue of Acr contains over 45 non homologous proteins [28]. Multiple large-scale genomics studies have shown that they often appear encoded on bacterial genomes within prophage regions [29]–[31]. More recently, a study of *P. aeruginosa* showed that accounting for the deactivation of CRISPR-Cas systems by co-located *acr* genes on the genome was linked to the genome size distribution [22]. Aside from CRISPR-Cas systems, bacterial genomes are homes to other lines of defense, which are often collectively referred to as the bacterial immune system. Historically, one of the first characterized set of genetic systems linked to phage defense are the restriction-modification systems, which are widespread throughout the bacterial domain [32]. The catalogue of defense systems has expanded over the years, with systems like BREX, CRISPR-Cas, Abortive infection (Abi), Toxin-Antitoxin, and DISARM [33]–[36]. Importantly, many new systems were discovered in recent years by leveraging the hypothesis that defense genes conglomerate in so-called defense islands [37]–[39]. The diversity of defense system options available to bacteria, for which these systems are typically found within the accessory genome content is now described as the pan-immune system [40].

In this study, we investigate the connections between the presence of CRISPR-Cas systems, anti-CRISPR proteins, defense system genes, genome size distributions, population structure, and phage susceptibility. We study these in *Pseudomonas aeruginosa*, a well-known and abundantly sequenced opportunistic and archetypal human pathogen, and its bacteriophages [41]. We show that (1) the population structure of this species has an impact on the distribution of genome sizes and CRISPR-Cas systems, (2) the anti-CRISPR proteins of type I-C and I-E appear to have a strong de-activation effect as proxied by genome sizes when compared to the type I-F, (3) the presence of CRISPR-Cas systems on a given strain from the population is positively correlated with phage susceptibility, and 4) the presence of CRISPR-Cas systems is linked to the depletion of other defense systems of the pan-immune system of *P. aeruginosa*.

## Materials and methods

### Bacterial strains and phage isolates

A total of 173 *Pseudomonas aeruginosa* isolates were collected from the glycerol stocks of three panels (Supplementary table ST1). All isolates were routinely cultured in Lysogeny Broth at 37 °C. The fourteen diverse and strictly virulent phages were retrieved from our stock (Supplementary table ST2).

### Host-range screening

Bacterial isolates were screened for their susceptibility to the phages using spotting dilution series assay [42]. Briefly, 10^2^ to 10^8^ dilution series of our phage stock were produced and up to 10 µl of each dilution were spotted on a double agar overlay. The plates were incubated overnight at 37 °C and checked the next day for signs of infection. We distinguish productive infection from other interactions through the plaque phenotype, i.e., whether individual plaques appear in the dilution series.

### Genome sequencing, assembly, and annotation

The genomic DNA of the isolates was sequenced using Illumina or MGI technologies (see Supplementary table S1). Raw sequencing files were controlled using FastQC and processed with Trimmomatic v0.40 for adapter clipping, quality trimming (LEADING:3 TRAILING:3 SLIDINGWINDOW:4:15), and filtering on length (>50 bp) [43]. Genomes were assembled using SPAdes v3.15.0 using the –careful option [44]. Quality of the resulting assemblies was assessed using Kraken 2 to assess potential contamination [45], QUAST v5.0.2 for assembly metrics [46], and the assembly graphs were visualized with Bandage v0.8.1 [47]. Additionally, some strains were re-sequenced on a MinION nanopore sequencer equipped with a R9.4.1 flowcell and a rapid preparation library (SQK-RBK004) (Supplementary table ST1, strains marked in italic). Long-read sequencing data were combined with short-read data for these strains and assembled with Unicycler v4.5.1 as detailed in [48].

### Genome datasets

Three datasets of genomes were analyzed in this study. The first one consisted of the 173 in-house assemblies described above and listed in Supplementary table ST1. The second dataset additionally includes 295 *P. aeruginosa* strains from the NCBI RefSeq database (accessed online 1/3/2021) that are marked as “Complete” assemblies [49] (Supplementary table ST3), for a total of 468 strains (173+295). Finally, the third dataset comprised 4,812 *P. aeruginosa* strains (Supplementary table ST4) obtained from the NCBI RefSeq database (release 98) that were further filtered based on genome length and GC% (accepted values: 5.7-7.5 Mbp and 65.4-67.0 respectively).

### Population structure

All genomes of the first dataset (173 strains) were annotated with Prokka v1.14.5 [50]. The annotated .gff files produced by prokka were analyzed using the pangenome software roary v3.13.0 using default options for the production of a core genome alignment [51]. The population phylogeny was established using iqtree v2.0.3 with 1,000 bootstrapping (UFBoot) and automatic model selection. The tree was visualized in iToL [52]. For this analysis, the dataset of 173 strains was complemented with 294 *P. aeruginosa* genomes marked as complete in the NCBI RefSeq database (Supplementary Table ST4). Those genomes were downloaded assembled and annotated, and the .gff files were obtained using the bioperl script Genbank_to_gff3.pl [53].

### Defense systems annotation

Protein accession numbers of *Pseudomonas aeruginosa* defense systems were retrieved from the PADS database [54] and obtained from the NCBI Protein database (Supplementary table ST5). We clustered proteins by sequence similarity using cd-hit v4.8.1 (75% identity over 75% length) to reduce the number of redundant entries for each system [55]. The proteome content of each of the 173 isolates was then iteratively queried using BLASTp against the (cd-hit) protein clusters of the different defense systems. The best hit was kept for each query, with requirements that the hit be of high significance and coverage (e-value < 10^-50, query coverage > 75%). CRISPR-Cas systems were separately annotated using CrisprCasTyper [56].

### Anti-CRISPR annotation

Acr proteins were annotated using the AcrFinder software [30] and we kept the homology hits (high confidence) and the Guilt-by-association (GBA) hits separated. We annotated 4,812 strains available in the NCBI RefSeq database with respect to their CRISPR-Cas type and anti-CRISPR (low confidence GBA hits were discarded). The strains were marked as CRISPR-when a co-located CRISPR-Cas and corresponding deactivating Acr were found, or CRISPR+ otherwise.

### Spacerome, genomic islands and prophages annotation

The spacers of the 173 strains equipped with CRISPR were aggregated and BLASTn (task: blastn-short) was used to check for their presence within the genomes of our phage collection (supplementary table ST2) and in the annotated defense system genes annotated in the 173 strains (requirement: identity with >90% coverage). The subset of 50 strains with closed genomes reconstructed with hybrid assembly was annotated with respect to their prophage and genomic island content using PHASTER and IslandViewer4 [57], [58]. For each strain, we calculated the intersection between the lists of defense system genes loci and the lists of genes included in the prophages and genomic islands separately (Supplementary table ST6).

## Results

### The population structure of *P. aeruginosa* and the distribution of CRISPR-Cas systems

We used a dataset of 468 sequenced strains and observed a bimodal distribution of genome lengths linked to the presence and absence of CRISPR-Cas systems as reported previously by Wheatley *et al*., 2021 (Supplementary figure 1 and supplementary table 3). In addition, we reconstructed the phylogeny of this dataset to account for the population group structure (Figure 1A). This revealed that the isolates from group II (n=178) have, on average, a larger genome than those from group I (n=293) (Figure 1B). Their average sizes differ by almost 200 kbp (6,643 kbp in group I and 6,842 kbp in group II), indicating that membership of isolates to these groups of the population structure of *P. aeruginosa* also informs the distribution of genome sizes. This analysis was not extended to the three other groups of the population (namely III, IV, and V with six, seven, and four isolates in the dataset, respectively), as they are currently under-represented in the public databases of *P. aeruginosa* genomes [59].

**Figure 1:**
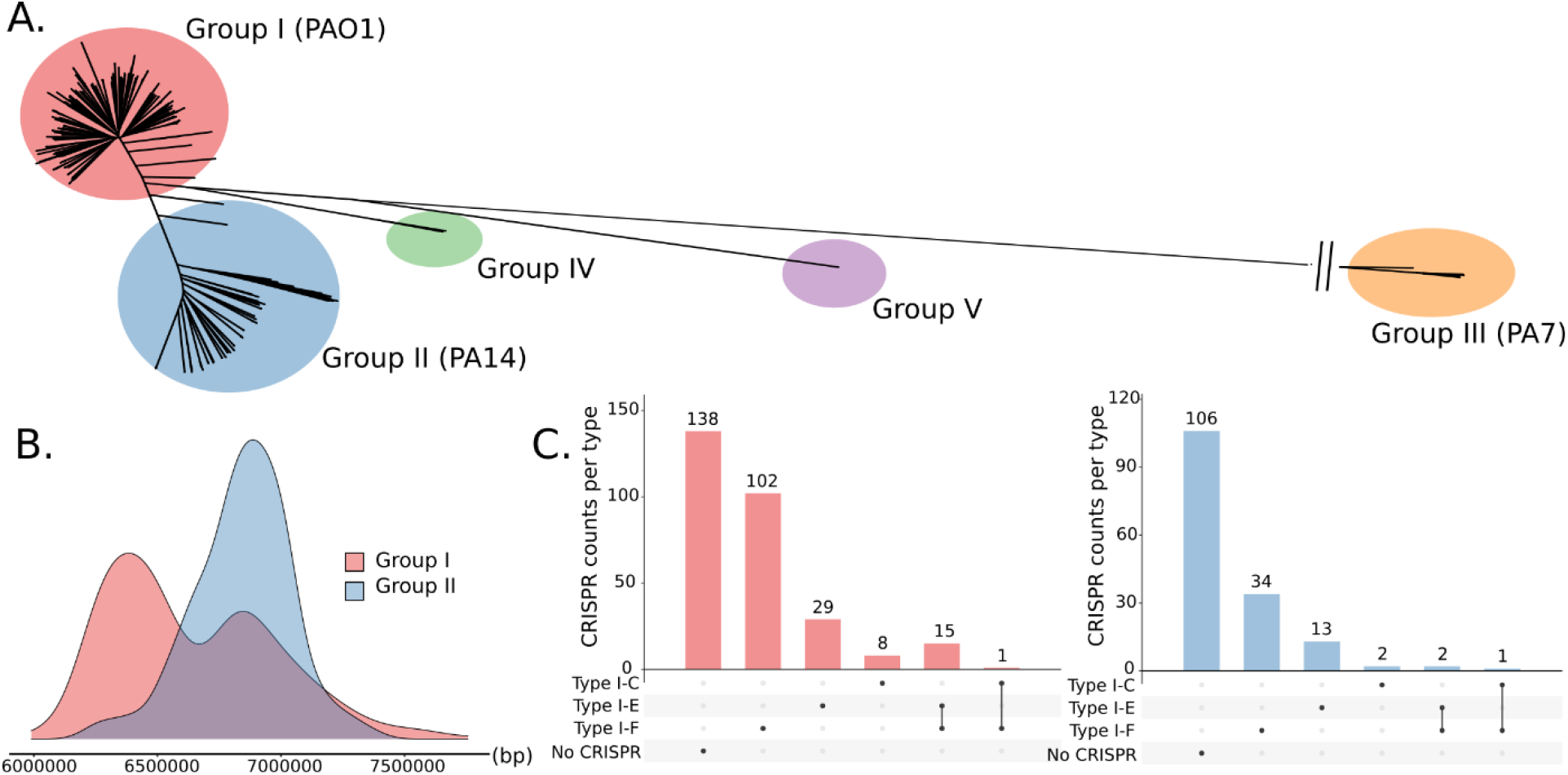
Association between the genome size of isolates and their position in the population structure. (A) The population structure of 468 genomes displays the typical five sub-groups population structure of P. aeruginosa. As few isolates originate from groups III, IV, and V, we omitted these groups from our analyses. (B) The genome sizes of isolates from group II are on average 200 kbp larger than those from group I, and seemingly do not display a bimodality of length distribution. (C) Tally of the CRISPR-Cas systems present in the groups I and II reveals that group II carry less (32%) CRISPR-Cas systems as those from group I (53%).

Interestingly, we also observed that the (on average larger) isolates of group II have a much lower propensity to carry a CRISPR-Cas system (Figure 1C), indicating that the population structure may be confounding the relationship between the presence of the CRISPR-Cas defense systems and their impact on the genome size. Indeed, about 53% of the isolates from group I appear to carry such a system, as opposed to 32% in group II. However, the bimodal distribution of genome sizes linked to CRISPR-Cas systems remained statistically significant within each group I and II (Wilcoxon test, p-value < 0.01). There is also a slight difference in the types of systems carried within these two groups. While in group I, the majority of strains with a CRISPR-Cas system have a type I-F system (75%), this number decreases to 69% in group II (Figure 1C).

### Accounting for anti-CRISPR deactivation of CRISPR-Cas systems I-C, I-E, and I-F

We annotated the RefSeq dataset with respect to the presence of CRISPR-Cas systems and anti-CRISPR (*acr*) genes found on the genomes (n=4,812 *P. aeruginosa* genomes, see Supplementary Table ST4) and found that 47% of the isolates were annotated with CRISPR-Cas systems (Figure 2A). This is in line with previous reports in *P. aeruginosa* [12], and some of these systems could be deactivated by the products of co-located *acr* genes (Figure 2B) [22]. When taking this deactivation into account, the effect on the genome size can be partially recapitulated in the systems I-C and I-E, but less so in type I-F systems (Figure 2C). This further counts towards the observation that isolates from group I (75% equipped with type I-F) are smaller on average than those of group II (69% equipped with type I-F) as the deactivation of those system appears to be more questionable.

**Figure 2.**
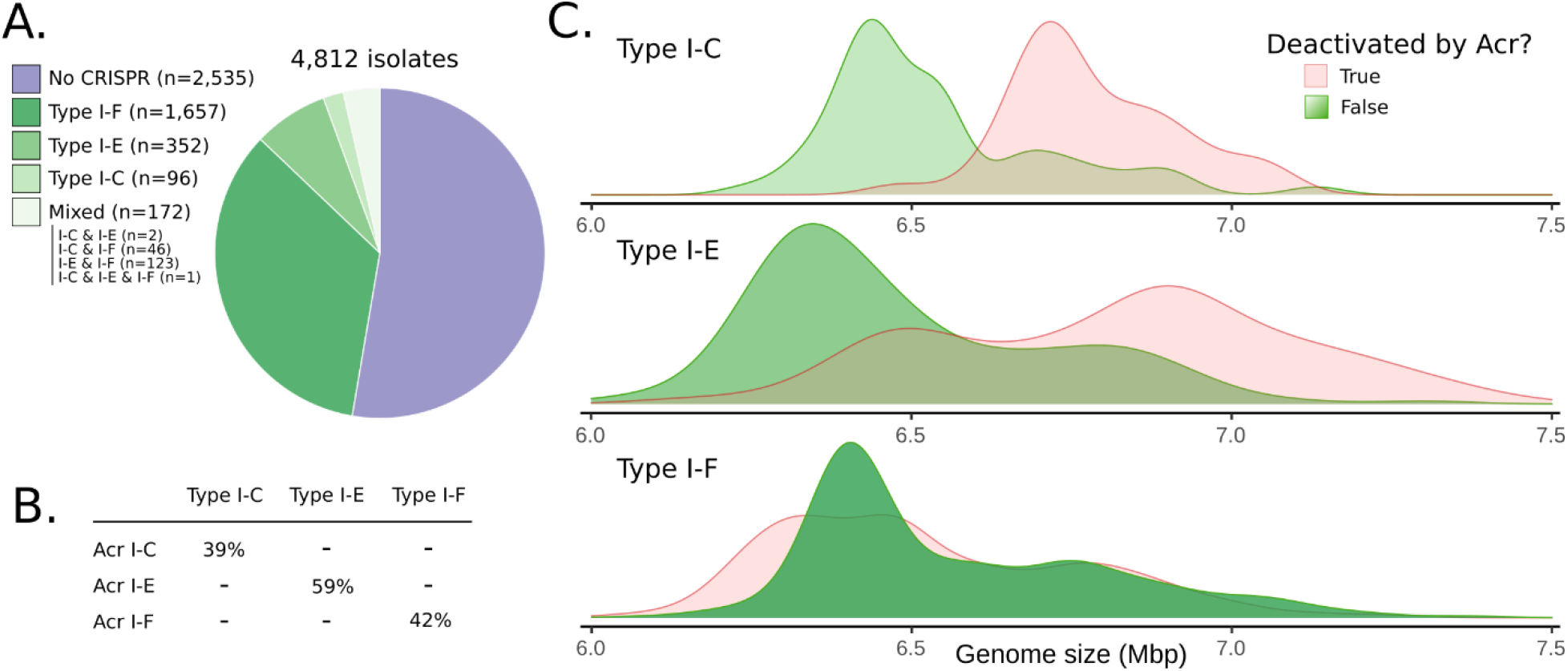
Deactivation of CRISPR-Cas systems by anti-CRISPR. (A) Distribution of the different CRISPR-Cas systems annotated in the RefSeq dataset from NCBI (n=4,812) shows that type I-F is prevalent (72%), followed by type I-E (15%), type I-C (4%), and various combinations of systems (7%). (B) Proportions of each system for which anti-CRISPR proteins are co-located on the genome. (C) Plots of genome sizes separated by types of CRISPR-Cas systems reveal that the inactivation of type I-F systems does not impact the distribution of genome sizes whereas anti-CRISPR deactivation of CRISPR-Cas systems of type I-C and type I-E is partially captured.

### Phage susceptibility of isolates is positively associated with the presence of CRISPR-Cas

We sequenced a collection of 173 *Pseudomonas aeruginosa* isolates that we screened against fourteen well-described virulent *Pseudomonas* phages from diverse genera [*Pbunavirus, Kmvvirus, Luz24virus, Elvirus, Phikzvirus, N4virus*] for signs of productive infection (single plaque phenotype) by spotting dilution series of the phages onto a double agar overlay (Supplementary table ST1 and ST2). These 173 *P. aeruginosa* genomes were annotated with respect to CRISPR-Cas, *acr* genes, and placed according to their group membership in the population structure (Supplementary Table ST7). In total, 55% of the strains were assigned to group I, 40% to group II and the remaining strains to group IV and V (no isolates from group III). We tabulated the overall susceptibility of these strains to the fourteen phages, as well as the susceptibilities of individual strains to the fourteen individual phage isolates (Supplementary table ST7 and ST8). Surprisingly, the strains whose genomes harbored CRISPR-Cas systems are on average more susceptible to productive phage infection (Figure 3A). This holds for individual phages (Figure 3B) and after accounting for deactivation of the system by Acr (CRISPR+/CRISPR-, supplementary figure 2). Finally, the effect is statistically significant when considering groups I and II separately (Wilcoxon test, p-value < 0.01).

**Figure 3:**
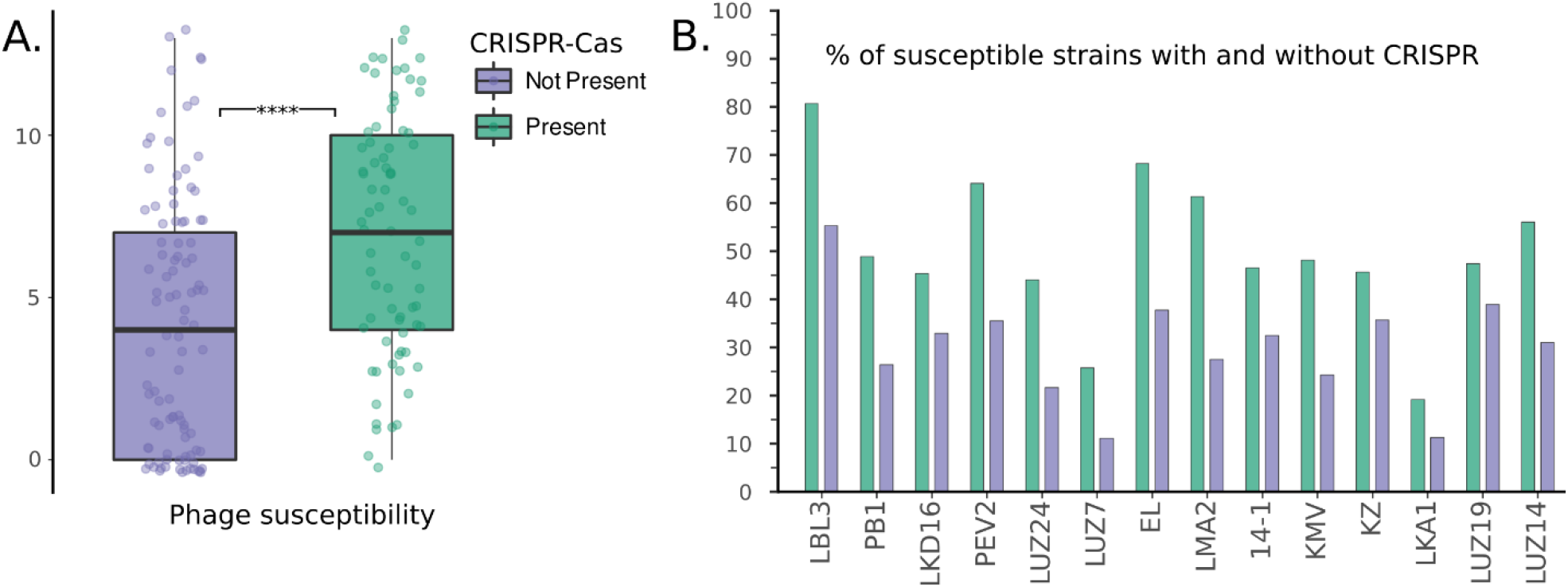
The impact of CRISPR-Cas on phage susceptibility. (A) The presence of CRISPR-Cas on bacterial genomes is correlated with a higher phage susceptibility. (B) This higher susceptibility is consistent across all 14 phages tested in our study.

In total, we found that 43% of the 173 strains carry a CRISPR-Cas system with 55% of group I strains, and 20% of group II strains (Figure 4A). The analysis of the spacers found in the CRISPR arrays of the strains revealed marginal amounts of hits towards the phages that we used, mostly against phage LMA2 although the few strains that possessed spacers against LMA2 appeared to be nevertheless susceptible to the phage with the strain PaLo170 being the only exception (Supplementary Table ST9).

**Figure 4:**
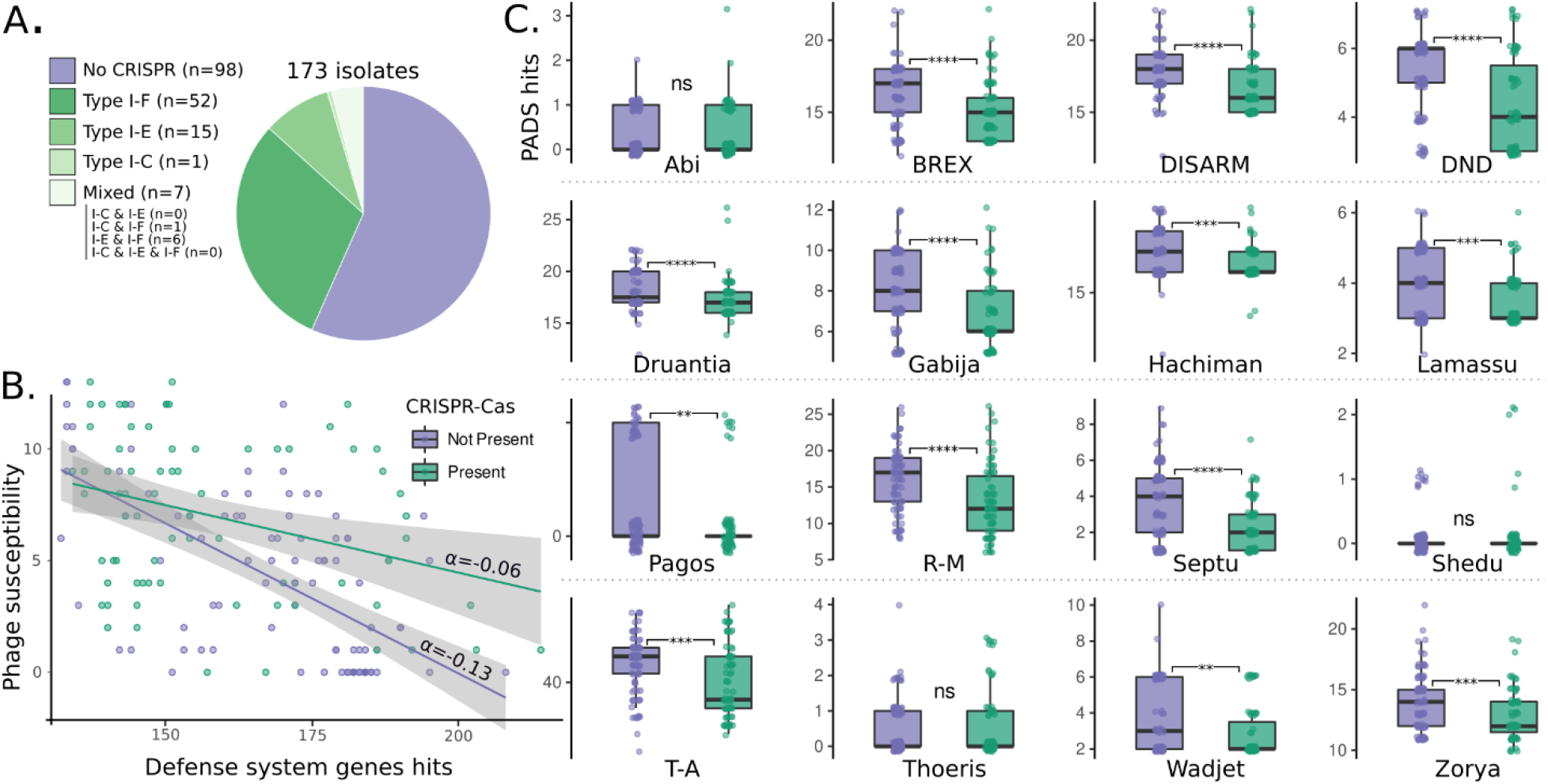
The impact of CRISPR-Cas on phage infectivity and defense system genes. (A) The proportion and types of CRISPR-Cas systems found in our dataset of 173 strains tested for their susceptibility follows that of the larger (sequenced) population (figure 2A). (B) Phage susceptibility versus the number of defense system genes annotated in each genomes, in strains with (α=-0.06) and without CRISPR-Cas (α=-0.13). (C) The number of genes present in 13 out of the 16 annotated defense systems versus the presence of CRISPR-Cas.

### Relationships between CRISPR-Cas and other defense system genes

We used the recently developed Prokaryotic Antiviral Defense Systems (PADS) database of proteins [54] to specifically annotate *P. aeruginosa* Defense System (DS) genes in the set of 173 strains and found a significant association between the number of accessory defense genes in the strains and phage infection (Figure 4B), as well as between the number of given defense system genes in strains with or without CRISPR-Cas systems (Figure 4C). Specifically, 13 out of the 16 annotated defense systems were significantly depleted when accounting for the presence or absence of CRISPR-Cas. The systems Abi, Shedu, and Thoeris did not show a statistically significant association with the CRISPR-Cas systems and were marginally detected in the dataset.

### Evidence of defense system genes targeting from spacers and their presence in genomic islands and prophages

The individual defense genes annotated in the dataset of 173 strains were queried against the spacers collected from those strains to find evidence of targeting of the defense system genes. Table 1 summarizes the hits found (>90% coverage) during the analysis. Most hit proteins are annotated as part of restriction-modification systems, and we can see that some spacers appear to be shared across multiple strains of the dataset. Interestingly, we also observe that some defense system genes are the target of multiple distinct spacers (e.g. as many as nine different spacers target loci of the gene which encodes the hypothetical protein KFL09433.1). In the results reported in the supplementary table ST10, we also highlight that spacer hits are mostly located in strains that lack a CRISPR-Cas system, as might be expected. We indicated in the last column of the Table 1 whether the defense genes are typically linked to prophage regions after manual inspection.

**Table 1.** Defense genes targeted by spacers: list of spacer hits in the defense genes of the 173 strains dataset. For each protein, we list the functional annotation, the defense system, the number of strains possessing a spacer against the gene coding for the protein, how many strains possess the target, how many distinct spacers there is, and whether the protein is linked to a prophage.

| Protein | Functional annotation | Defense System | #Strains w/ spacer | #Target Strains | #Different spacers? | Linked to Prophage? |
| --- | --- | --- | --- | --- | --- | --- |
| **KFL09433.1** | Hypothetical protein | R-M | 31 | 6 | 9 | Yes |
| **OPA45023.1** | Cytosine methyltransferase | R-M | 15 | 57 | 6 | Yes |
| **OKR21085.1** | Hypothetical protein | R-M | 4 | 17 | 3 | Yes |
| **OZO08709.1** | Conjugative transfer ATPase | DND | 12 | 121 | 3 | No |
| **KWX42418.1** | DNA Methyltransferase | R-M | 2 | 8 | 2 | No |
| **ABR80919.1** | Adenine methyltransferase | R-M | 3 | 11 | 1 | Yes |
| **AHH52652.1** | Cytosine methyltransferase | R-M | 3 | 9 | 1 | Yes |
| **AID83532.1** | HNH Endonuclease | Septu | 3 | 34 | 1 | Yes |
| **EMZ50108.1** | Hypothetical protein | R-M | 1 | 1 | 1 | Yes |
| **KFL12158.1** | Phosphoadenosine reductase | DND | 8 | 14 | 1 | Yes |
| **KSF89549.1** | Cytosine methyltransferase | R-M | 4 | 14 | 1 | No |
| **OTI54595.1** | SNF2 family helicase | DISARM | 1 | 95 | 1 | No |
| **OZO07714.1** | DEAD/DEAH box helicase | BREX | 1 | 48 | 1 | No |

Finally, we analyzed the presence of prophages and genomic islands in a subset of 50 strains that we re-sequenced with nanopore sequencing and assembled into complete sets of replicons. In that subset of strains, we looked at the location of the annotated defense system genes and whether these locations coincide with either prophages or genomic islands (from which we excluded prophages detected to keep the two contributions clearly separated). In figure 5, we plot the count of genes that are found in either prophage regions (Figure 5A) or genomic islands (Figure 5B) and their contribution to the total number of defense system genes. We note that defense system genes within prophage regions contribute only slightly to the total amount of defense genes (Pearson correlation coefficient: 0.3). On the other hand, genes associated with genomic islands contribute the majority of defense genes in any given strain (Pearson correlation coefficient: 0.93).

**Figure 5:**
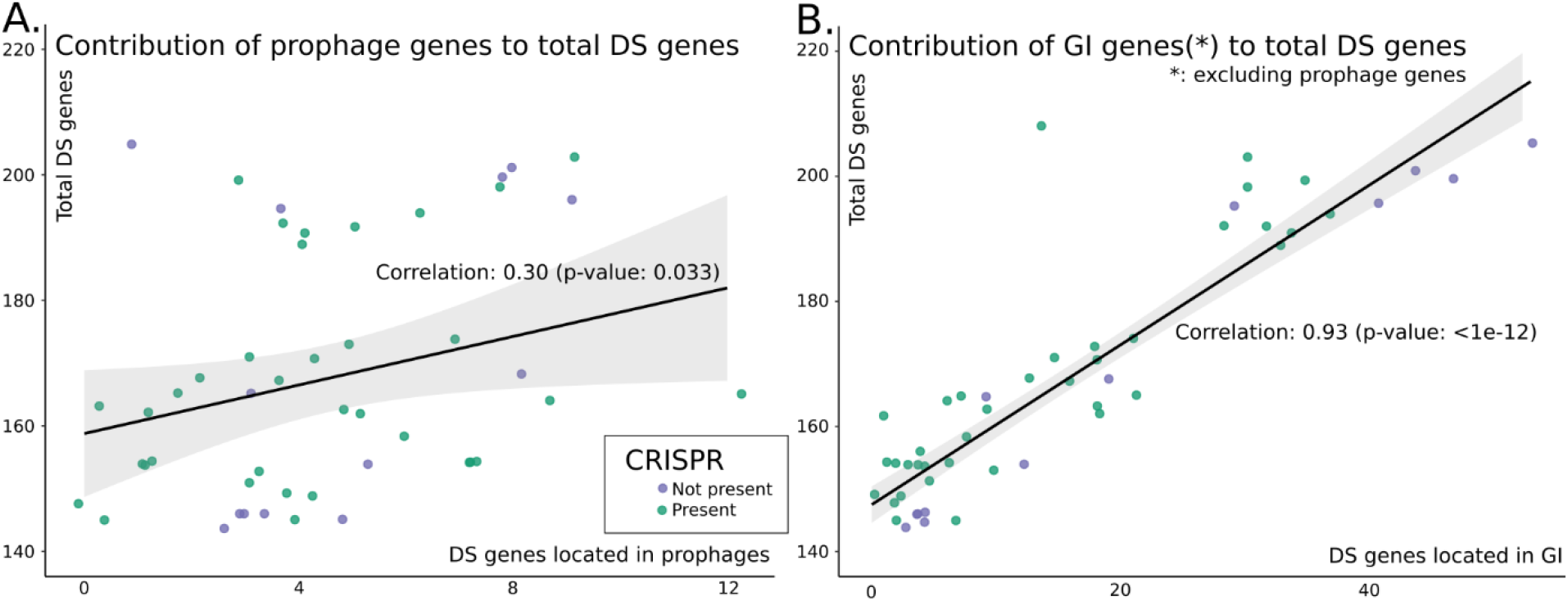
Contributions to the total number of defense system genes. (A) from annotated prophage regions (B) from genomic island regions from which we removed intersection with prophage regions.

## Discussion

### The distribution of P. aeruginosa genome sizes is linked with the presence of CRISPR-Cas, the population structure, and anti-CRISPR activity

Wheatley *et al*. [22] recently highlighted an association between smaller genome size and the presence of CRISPR-Cas in 300 *P. aeruginosa* genomes. The authors proposed that the xenogenic nature of CRISPR-Cas works as a general mechanism constraining horizontal gene transfer (HGT), thereby preventing the acquisition of new genetic material, typically accompanied with a gain in genome size and a decrease in GC% in that species [60]. Here we propose that additional information can be gathered by including the population structure information of that species into account [61]. Indeed, the population structure of *P. aeruginosa* may be considered in five groups, with groups I and II containing the vast majority of the isolates sequenced to date [59]. We show that the distribution in genome size is strongly linked with the group structure and predicts to some degree the presence of CRISPR-Cas systems, which are present in about half of group I isolates (53%) and a third of group II (32%). Importantly, although the group structure appears is confounding the relationship between the presence of CRISPR and the genome size, our data support the original hypothesis of the overall impact of CRISPR-Cas on HGT, including in both sub-groups. This impact of CRISPR-Cas systems on population evolution within a given bacterial clade was previously reported in the *Bacillus cereus* group[21], within which the deactivation of CRISPR-Cas systems of the different species in that group appeared to correlate with wider environmental distribution. We show here that within a single species, *P. aeruginosa*, this general mechanism may also be at work.

We also investigated the link between the occurrence of co-located *acr* genes on CRISPR-Cas deactivation of and the impact on genome size. As some types are less represented (I-E and I-C), we used the larger RefSeq dataset consisting of 4,812 sequenced strains to show a significant association between the sizes of genomes equipped with CRISPR-Cas systems and those in which the system is present but where it appears to be deactivated by co-located antiCRISPR proteins [35], [36]. This hypothesis was previously reported for *P. aeruginosa*, but we show here that the activity of antiCRISPR differentially affects the three different systems typically detected within the species, i.e. type I-C, I-E, and I-F [11]. Indeed, we see that the distribution of genome sizes can be partially recapitulated by the deactivation of CRISPR-Cas systems type I-C and I-E. Interestingly, while type I-F appears mostly unaffected by the activity of Acr proteins (Figure 2C), despite the previously demonstrated efficacy of Acr type I-F proteins against CRISPR-Cas systems I-F in a strain from that very same species [25]. Taken together with the link between the population structure, genome sizes, and CRISPR-Cas, our data indicate that these factors should be explicitly considered when running large computational analysis of the impact of CRISPR-Cas systems on the genome of a given species.

### CRISPR-Cas may prevent the acquisition of other effective phage resistance systems and result in higher susceptibility to virulent phages

We showed an increased overall phage susceptibility in strains which harbor CRISPR-Cas systems in their genomes. This apparent paradox, given the established role of CRISPR-Cas as a defense system against phage infection [62], [63] holds for all 14 phages that we studied and after accounting for population structure. Others have proposed that CRISPR-Cas may not be the principal defense mechanism behind the resistance to phages in multi-phage resistant strains of *P. aeruginosa*, highlighting the absence of CRISPR-Cas in phage-resistant strains [64], [65]. We show here that the effect is not limited to multi-phage resistant strains, but in fact occurs across the panel and influences overall susceptibility of the population.

To further understand what may be driving this increased susceptibility, we considered the accessory genome components of the pan-immune system that have been previously linked to phage resistance [40]. Indeed, CRISPR-Cas systems are but one defense mechanism in bacteria, with many others described in *Pseudomonas* [33], [36], [38], [66]. In the presence of CRISPR-Cas, all but three of the 16 pan-immune systems we specifically considered in our analysis are depleted. This would suggest that while CRISPR-Cas based adaptive immunity may provide protection to individuals in a CRISPR-enabled population invaded by horizontal elements including bacteriophages, the population may have traded off the capacity for the acquisition of other, non-adaptive and perhaps more valuable, phage defense systems. Additionally, we see a strong correlation between the number of defense system genes and overall susceptibility to phages. The decrease in susceptibility appears to be true regardless of the presence of CRISPR-Cas but seems stronger in the strains with CRISPR-Cas (Fig 3-B). It is tempting to speculate that this could be the result of decreased HGT, which then limit the organisms’ capacity for adaptation in dynamic environments, or a crosstalk between the defense systems in preventing phage infection, which CRISPR-Cas could be impeding. Importantly, further research will be necessary to tease apart these dynamics as new defense systems are likely still to be discovered amongst the bacterial genomes, their annotation in the genome may not necessarily correlate with their functioning or depend on specific environmental conditions, and the regulation of the expression of CRISPR-Cas and these other systems remains under investigation [67], [68].

### Is there evidence that the pan-immune system is a target of the CRISPR-Cas system?

We looked for evidence of targeting of phage defense systems given the negative association with CRISPR-Cas, either by evidence of spacers that could target defense genes directly, or indirectly via the targeting of genomic islands. In the first case, we found that some genes discovered during our search were the target of multiple distinct spacers, with a currently uncharacterized protein having as many as nine distinct spacers against it (Table 1). Many of the genes targeted belong to the restriction-modification systems, and upon manual inspection appeared to be within the family of methyltransferases, known for their role in R-M systems which protect against phages [69]. A subset of genes detected are associated with prophage regions, and as such it is difficult to discriminate whether they are the target of spacers because of their function or because they are located on mobile prophage genomes. We also found that the genes targeted by spacers are mostly located on strains that do not possess a CRISPR-Cas system, in line with our expectations of the functioning of CRISPR-Cas and our results showing a negative association with the other defense systems.

Additionally, we re-sequenced 50 strains from our panel with long read technology, using hybrid assembly to assemble closed replicons [70]. This enabled us to annotate the presence of prophages and other genomic islands amongst the genomes and avoid the problems of concatenating contigs or re-ordering them using a reference genome during their detection in draft genomes [71]. By intersecting for each of these strains the list of defense genes and their position along the genome with the loci of the prophages on the one hand, and the genomic islands on the other, we showed that prophage contribute some defense system genes to the total pool of defense system genes. However, this correlation appears relatively limited (Pearson correlation coefficient: 0.3). On the other hand, the genomic islands appear to include many defense system genes, and these contribute to a large degree to the total count of defense system genes in the strains we analyzed (Pearson correlation coefficient: 0.93). This result is consistent with previously published observations on the effect of CRISPR-Cas on evolutionary dynamics in *P. aeruginosa* [22]. As defense genes have been shown to be co-located on so-called defense islands [37], [38] it appears that the depletion in prophages and ICEs observed in *P. aeruginosa* could result in a limitation of defense systems available in the isolates equipped with CRISPR-Cas. An important corollary of this is that antibiotic pressure, which is classically driving selection of MGEs, may mitigate against CRISPR-Cas. One may wonder then if antimicrobial resistance and phage resistance will be co-selected.

## Concluding remarks

We showed here what appears at first to be a paradoxical link between the increased phage susceptibility of *P. aeruginosa* strains and the presence of CRISPR-Cas systems, known for their potential to prevent phage infections. To understand what could be driving this phenomenon, we investigated whether a trade-off exists between these systems and other characterized anti-phage defense systems found in the pan-immune system of bacteria. This allowed us to show a depletion of defense genes on the isolates of the population that carry CRISPR-Cas. We also found evidence of targeting of such defense genes either directly by analyzing CRISPR array targets, and via their location on genomic islands. This association with genomic islands suggests that important phage resistance is commonly horizontally acquired, as is the case for resistance to antimicrobial drugs. Our finding of a negative correlation between phage resistance and CRISPR-Cas systems, which effectively defend against HGT, may be less paradoxical when seen in that light. It could reflect the importance of the other components of the pan-immune to protect against phage infection and the evolutionary role of CRISPR-Cas systems. It also opens the door to further investigation regarding the potential of this dynamic to co-select antimicrobial and phage resistance.

## Data availability Statement

All sequencing data was deposited in the NCBI SRA database and is accessible via the BioProject PRJNA720041.

## Acknowledgements

CL is supported by a PhD fellowship from FWO Vlaanderen (1S64720N). The labs of VvN and RL are supported by the ID-N grant PHAGEFORCE from the KU Leuven (IDN/20/024). The authors thank Frédéric Fux and Alison Kerremans for excellent technical assistance. We would also like to thank team at Westmead Institute for Medical Research in Sydney, specifically Ruby Lin for sequencing support of 60 P. aeruginosa isolates (MGI/Decode Science/Micromon Genomics Next Generation Sequencing Research Grant 2019 AU20190724-02), Nouri Ben Zakour for help in selection of strains for testing and Matthew Smith for excellent laboratory assistance.

## Supplementary figures

**Supplementary figure 1:**
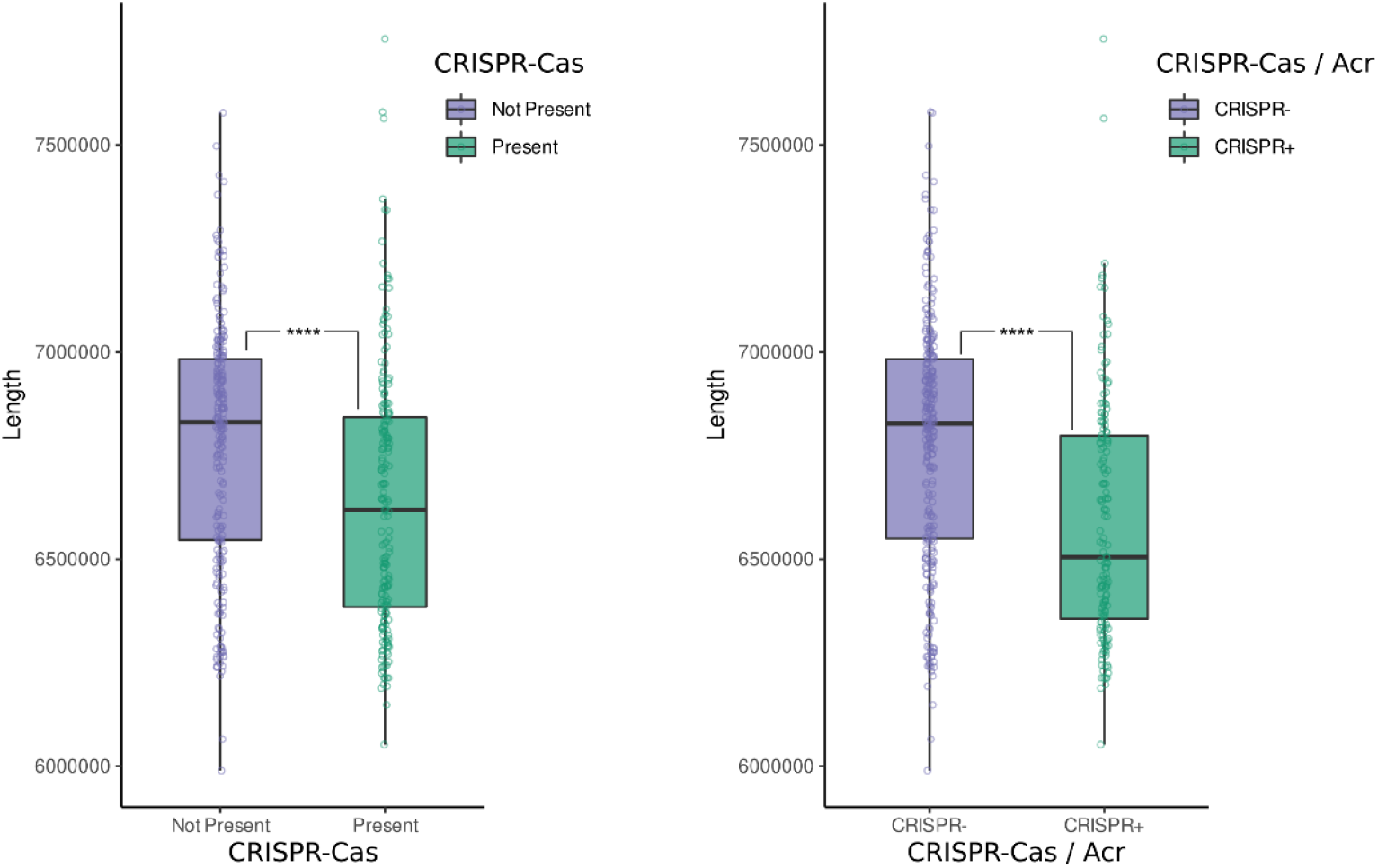
The left-hand side panel highlights the distribution of genome lengths with respect to the presence or absence of CRISPR-Cas systems. The right-hand side panel highlights the same, but accounts for potential de-activation of the systems by co-located Acr genes (CRISPR-)

**Supplementary figure 2:**
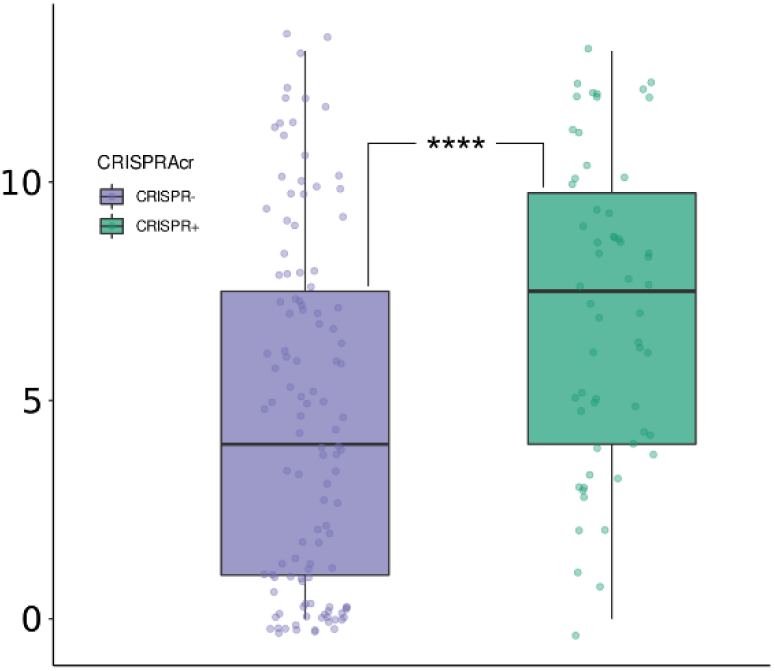
The presence joint presence of the CRISPR-Cas system and co-located *acr* genes on bacterial genomes is correlated with phage susceptibility (CRISPR-strains are less susceptible to phages).

## Supplementary tables

**Supplementary table ST1**: List of the 173 bacterial isolates sequenced for this study and screened for their phage susceptibility with their corresponding NCBI accession numbers. The strains re-sequenced with nanopore (n=50) are indicated in italic.

**Supplementary table ST2:** List of the fourteen virulent phages used in this study, with the accession number of their respective genomes and antiCRISPR annotation.

**Supplementary table ST3:** CRISPR-Cas and antiCRISPR annotation of 294 genomes available in the NCBI RefSeq database with assemblies marked as “Complete”, supplemented with the 174 genomes of this study (ST1).

**Supplementary table ST4:** Details of the isolates used for the CRISPR-Cas and antiCRISPR annotation (RefSeq dataset release 98, n=4,812)

**Supplementary table ST5:** List of proteins from the different defense systems of *Pseudomonas aeruginosa* retrieved from the PADS database [4]

**Supplementary table ST6:** Table with the annotation of the dataset of 173 strains, which includes the various annotations used in this study, including the defense systems, population structure group, Acr, phage susceptibility (0 = no infection, 1 = lysis effect, 2 = plaques observed).

**Supplementary table ST7:** Breakdown for each phage of the host-range results. Top-left: CRISPR-Cas presence only (not accounting for Acr activity) and only considering productive infections (individual plaques). Top-right: same, but with respect to Acr (CRISPR+/CRISPR-). Bottom-left: CRISPR-Cas presence only (no Acr), and considering both lysis effects and productive infections. Bottom-right: same, with with respect to Acr (CRISPR+/CRISPR-)

**Supplementary table ST8:** List of all spacer hits found in the collection of 173 strains that have targets against the phages used in our study.

**Supplementary table ST9:** Results of the intersection between prophage regions and genomic islands and the defense system genes discovered in each strain.

**Supplementary table ST10:** List of all spacers found in the dataset of 173 strains that had direct targets in annotated defense systems genes. The dataset has 43.4% of strains with CRISPR-Cas systems, and we have colored in green when the genes targeted are mostly found in strains that lack CRISPR systems, and in orange otherwise.

